# A better way to define and describe Morlet wavelets for time-frequency analysis

**DOI:** 10.1101/397182

**Authors:** Michael X Cohen

## Abstract

Morlet wavelets are frequently used for time-frequency analysis of non-stationary time series data, such as neuroelectrical signals recorded from the brain. The crucial parameter of Morlet wavelets is the width of the Gaussian that tapers the sine wave. This width parameter controls the trade-off between temporal precision and frequency precision. It is typically defined as the “number of cycles,” but this parameter is opaque, and often leads to uncertainty and suboptimal analysis choices, as well as being difficult to interpret and evaluate. The purpose of this paper is to present alternative formulations of Morlet wavelets in time and in frequency that allow parameterizing the wavelets directly in terms of the desired temporal and spectral smoothing (as full-width at half-maximum). This formulation provides clarity on an important data analysis parameter, and should facilitate proper analyses, reporting, and interpretation of results. MATLAB code is provided.

## Motivation for time-frequency analysis

Many biological and physical systems exhibit rhythmic processes. Rhythmic temporal structure embedded in time series data can be extracted and quantified using the Fourier transform or other spectral analysis methods.

However, the Fourier transform has a “soft assumption” of signal stationarity, which means that the spectral and other features of the signal remain constant over time. To be sure, the Fourier transform is a perfect representation of the time series, regardless of its temporal dynamics. However, the power spectrum resulting from the Fourier transform is easily visually interpretable only for stationary signals; non-stationarities are encoded in the phase spectrum, which is typically impossible to interpret visually.

Signal non-stationarities are the primary motivation for time-frequency analyses, in which the power spectrum is computed over short windows of time. The primary assumption here is that the signal is roughly stationary over some shorter (sliding) time window. For neural time series data, this assumption is typically a few hundreds of milliseconds. An example of a “static” and “dynamic” spectral representation of a non-stationary signal is presented in Figure 1.

**Figure 1.**
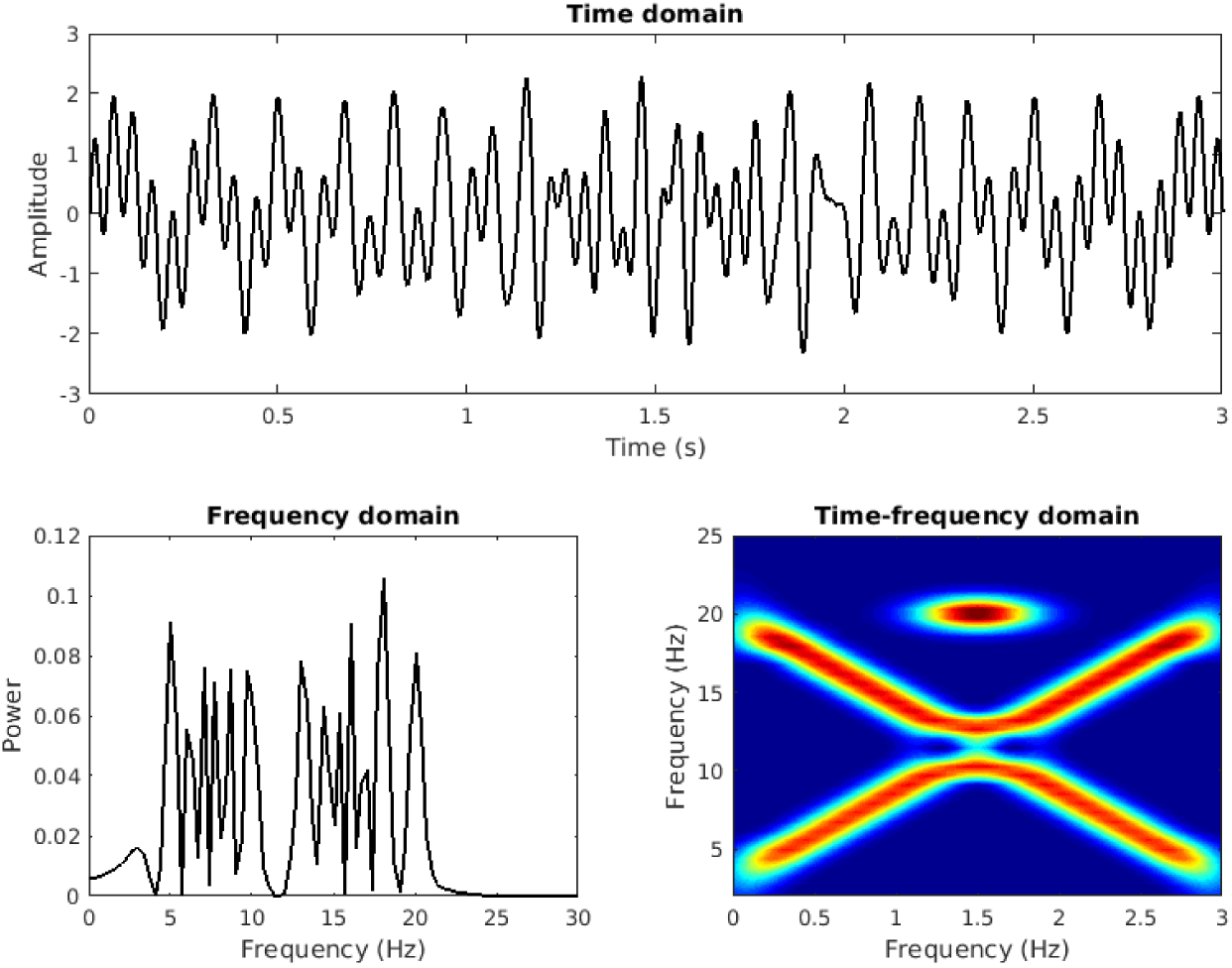
In the presence of signal non-stationarities (panel A), the “static” power spectrum from the Fourier transform can be difficult to interpret (panel B). In these cases, a time-frequency analysis (panel C) is often insightful.

## Overview of methods for time-frequency analysis

There are several time-frequency analysis methods, most of which produce qualitatively or quantitatively similar results (Bruns 2004; Cohen 2014). The three most commonly used are the short-time fast Fourier transform, complex wavelet convolution, and filter-Hilbert. These three methods have in common that they involve isolating a narrow temporal window of data and then extracting its frequency spectrum.

A key parameter in time-frequency analysis is the one that governs the trade-off between temporal precision and spectral precision; it is not possible to have simultaneously arbitrarily good precision in both time and in frequency. Furthermore, the resulting smoothness is often advantageous for averaging data across stimulus repetitions and individuals; thus, arbitrarily good precision can be detrimental in applied data analysis of biological systems.

*The key argument of this paper* is that this crucial parameter is created and reported in a way that obscures both the signal-processing and the theoretical assumptions that are imposed on the data, and that shape the results. *The key upshot of this paper* is two alternative methods for creating and reporting Morlet wavelets in a way that makes this key parameter transparent and easily interpretable.

## Advantages and assumptions of Morlet wavelets for time-frequency analysis

A Morlet wavelet is defined as a sine wave tapered by a Gaussian (Figure 2, top row). For time-frequency analysis, a complex Morlet wavelet is used, in which the Gaussian tapers a complex sine wave. The complex Morlet wavelet is then convolved with the time series signal, and the result of convolution is a complex-valued signal from which instantaneous power and phase can be extracted at each time point. Wavelet convolution can be conceptualized as a “template-matching” procedure, in which each time point in the signal is compared against a template (the Gaussian-windowed sine wave), and the result of the convolution is a time series of “similarities” between the signal and the wavelet.

**Figure 2.**
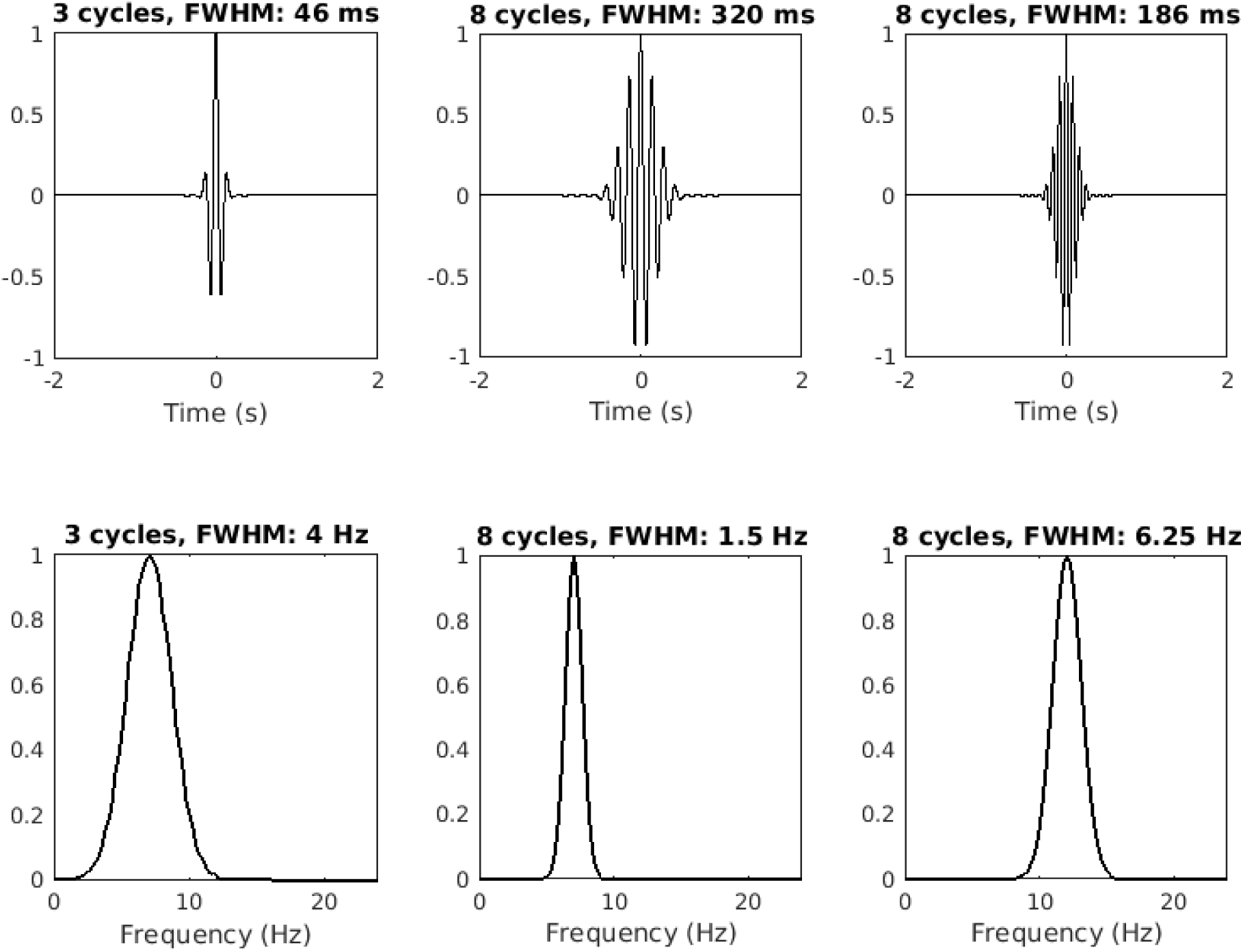
Three Morlet wavelets in the time domain (upper row) and in the frequency domain (lower row) with two frequencies and different time-frequency trade-off parameters. The argument of this paper is that reporting this parameter in terms of full-width at half-maximum (FWHM) in the time and/or frequency domains is more informative than number of cycles.

The tapering Gaussian has one parameter that defines its width (also called its shape or deviation). A wider Gaussian leads to decreased temporal precision but increased spectral precision, and vice-versa for a narrower Gaussian. This parameter is typically defined as the “number of cycles,” but the purpose of this paper is to argue that it would be better to define the Gaussian width as the full-width at half-maximum (FWHM), which is the distance in time between 50% gain before the peak to 50% gain after the peak.

There are several advantages of Morlet wavelets for time-frequency analysis. One is that the Morlet wavelet is Gaussian-shaped in the frequency domain (Figure 2, bottom row). The absence of sharp edges minimizes ripple effects that can be misinterpreted as oscillations (this is a potential danger associated with plateau-shaped filters). Second, the results of Morlet wavelet convolution retain the temporal resolution of the original signal. Third, wavelet convolution is more computationally efficient and requires less code compared to other methods, because it involves the smallest number of computations, most of which are implemented using the fast Fourier transform.

There are two key assumptions of Morlet wavelet-based time-frequency analysis. The first is that the signal is roughly sinusoidal. Some researchers have noted that brain oscillations are probably not shaped like pure sine waves (Cole and Voytek 2017; Jones 2016). On the other hand, the overwhelming success and ubiquity of time-frequency analysis methods indicates that minor violations of this assumption are not deleterious for practical data analysis, hypothesis-testing, data mining, and feature discovery. The sinusoidal assumption is therefore considered valid, and additional detailed waveform analyses can be insightful in specific instances.

The second assumption of Morlet wavelet analysis concerns stationarity of the signal. Interpreting the results of Morlet wavelet convolution relies on the assumption that the signal is stationary within the time window that the wavelet has non-zero energy. Generally, this can be taken as the FWHM of the Gaussian that is used to create the Morlet wavelet. Any signal non-stationarities that are smaller than this FWHM will produce results that are difficult to interpret (similar to the “static” power spectrum in Figure 1).

Appropriate selection of the width of the wavelet is a combination of signal processing considerations, and theoretical/speculative considerations of the system from which the signal was measured. This parameter is therefore important for time frequency data analysis, and yet is often selected and reported in a way that obscures the assumption underlying the data analysis. The key goal of this paper is to present two different methods of formulating and describing Morlet wavelets in a way that makes this assumption transparent and easily interpretable.

## Typical definition of Morlet wavelets

A complex Morlet wavelet *w* can be defined as the product of a complex sine wave and a Gaussian window:

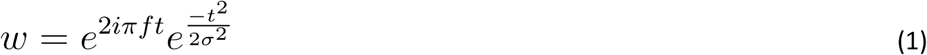

where *i* is the imaginary operator 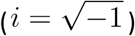, *f* is frequency in Hz, and *t* is time in seconds. To avoid introducing a phase shift, *t* should be centered at *t=0*, for example, by defining *t* from −2 to +2 seconds. σ is the width of the Gaussian, which is defined as

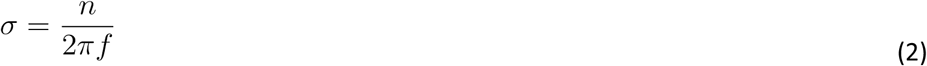

Bhe parameter that defines the time-frequency precision trade-off is *n*, which is often referred to as the “number of cycles.” For neurophysiology data such as EEG, MEG, and LFP, typical values of *n* range from 2 to 15 over frequencies between 2 Hz and 80 Hz. Other applications may require a different number of cycles.

There is nothing “wrong” with equations 1 and 2. However, the relationship between the number of cycles and the temporal-spectral smoothing is unclear, partly because most people do not think about time in terms of “number of cycles,” and partly because the effect of *n* on the temporal smoothing depends on the frequency of the wavelet (see Figure 2). Many researchers and students struggle with how to specify and interpret this parameter, which leads to confusion and possibly suboptimal data analyses.

## A better definition of the Morlet wavelet

A definition of the complex Morlet wavelet that provides more clarity is

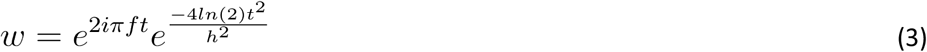

The new parameter here is *h*, which is the FWHM in seconds.

Equation 3 is not fundamentally different from equations 1-2. It is simply a way of rewriting the formula to make the temporal smoothing parameter easier to define, understand, and report. Rather than specifying a number of cycles (which might feel arbitrary and mysterious), one can specify a FWHM in seconds (assuming that *t* is defined in seconds), which is a more natural unit to think about in terms of desired temporal smoothing, and in terms of the assumption of the duration of stationarity in the system under investigation.

The *h* parameter in equation 3 is analytical; its actual implemented value depends on the data sampling rate. For this reason, it is recommended to compute and examine the empirical FWHM that is implemented in software. When the sampling rate is considerably higher than the frequency of interest, the specified and empirical widths should match closely.

The empirical FWHM can be obtained by subtracting the time point where the post-peak Gaussian is closest to 50% gain (if the Gaussian is normalized to a peak amplitude of 1, then this is the value closest to.5) from the time point where the pre-peak Gaussian is closest to 50% gain. (This algorithm is presented in the MATLAB code at the end of this document.) Figure 3 shows the relationship between the empirical FWHM derived from equations 1-2 vs. equation 3.

**Figure 3.**
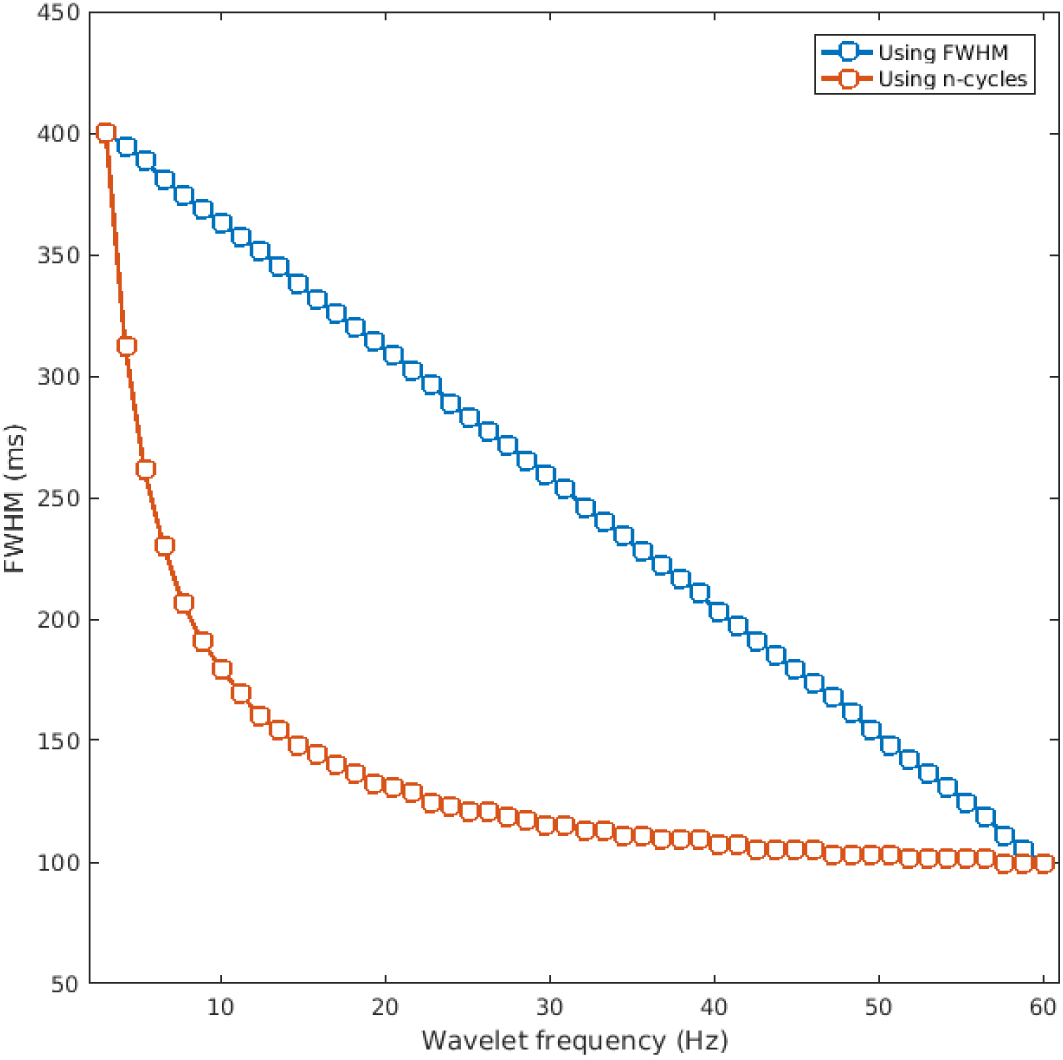
The empirical FWHM of the Gaussian (y-axis) used to define complex Morlet wavelets over a range of frequencies (x-axis), defined using equations 1-2 vs. equation 3.

## Recommendation for minimum wavelet FWHM

As mentioned earlier, there are two considerations for selecting the width of a wavelet for time-frequency analysis: signal processing and system quasi-stationary. The minimum width of a wavelet should be derived only from signal processing considerations.

It is recommended to set the FWHM no lower than one cycle at the frequency of the sine wave used to create the wavelet. For example, a 10 Hz wavelet should not have less than a 100 millisecond FWHM. This leads to the simple formula to compute the minimum FWHM of *1/f*, where *f* is the frequency of the sine wave that creates the wavelet. This corresponds to ˜2⅔ cycles (the *n* parameter in equation 2).

Note that a FWHM corresponding to one cycle actually means that more than one cycle will contribute to the wavelet, because there is still non-zero energy beyond the 50% gain boundaries used to define FWHM.

Also note that this is a recommendation for the *minimum* wavelet length. It is often sensible to have wavelets longer than the minimum, depending on a desired amount of temporal or spectral smoothing. One should also keep in mind that wavelets with narrower Gaussians (smaller FWHM) have wider spectral energy windows (Cohen 2014), so creating wavelets using the minimum bound is not necessarily optimal.

## Alternative: defining the wavelet in the frequency domain

The discussion above concerns defining the wavelet in the time domain. This is a natural approach in many situations, but in other situations it might be more natural to start from a desired amount of spectral smoothing. Thus, one can specify the desired spectral smoothing, which means defining the wavelet in the frequency domain instead of in the time domain. The formula is slightly different.

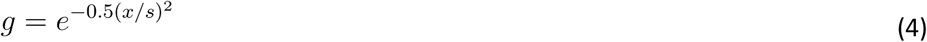

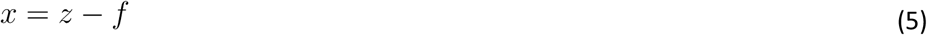

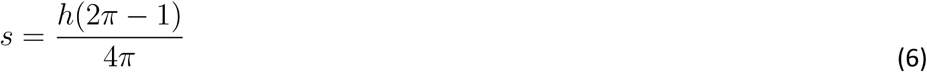

where *z* is a vector of frequencies in Hz from 0 to the sampling rate in the number of steps as the desired length of the spectrum (e.g., the length of the time series signal, or the length of the result of convolution), *f* is the peak frequency of the wavelet, and *h* is the FWHM in Hz.

The Gaussian in equation 4 creates the amplitude shape in the frequency domain; the time-domain wavelet can be obtained by the inverse Fourier transform of this envelope. The empirical FWHM in the time domain Morlet wavelet can be obtained by applying the FWHM estimation procedure to the magnitude of the complex time series. An example is shown in Figure 4.

**Figure 4.**
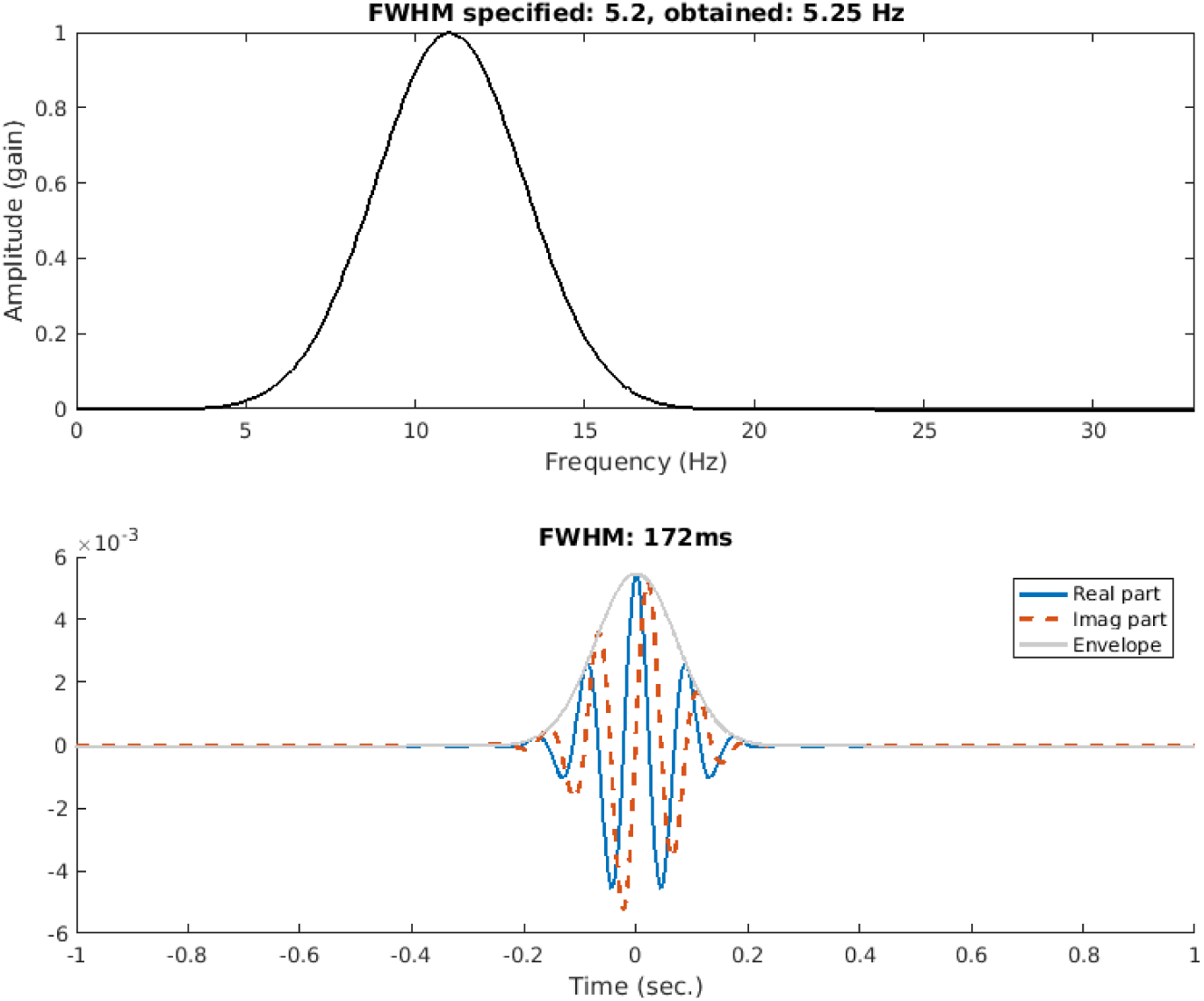
An example of specifying the wavelet shape in the frequency domain (top panel) and computing its inverse Fourier transform to obtain a time-domain Morlet wavelet (bottom panel). The temporal FWHM is defined using the envelope.

## Reporting

It is sensible and useful to report the FWHM in both the time and the frequency domains. This will facilitate interpretation of the results and replication of analysis methods. Here is a suggestion for describing the wavelets:

> We implemented time-frequency analysis by convolving the signal with a set of complex Morlet wavelets, defined as complex sine waves tapered by a Gaussian. The frequencies of the wavelets ranged from X Hz to X Hz in X steps. The full-width at half-maximum (FWHM) ranged from XXX to XXX ms with increasing wavelet peak frequency. This resulted in a spectral FWHM range of XX Hz to XX Hz.

## Conclusion

The goal of this paper is to provide an alternative way to conceptualize, define, and report Morlet wavelets for time-frequency analysis. No new methods, formulas, or algorithms are presented here; instead, the idea is to encourage researchers and students to make a minor adjustment to their existing analysis pipeline in a way that makes the findings, analysis methods, assumptions, and results, easier to understand, communicate, and interpret. MATLAB code to reproduce the figures is presented as a supplement.

---

**MATLAB code to reproduce figure 1.**
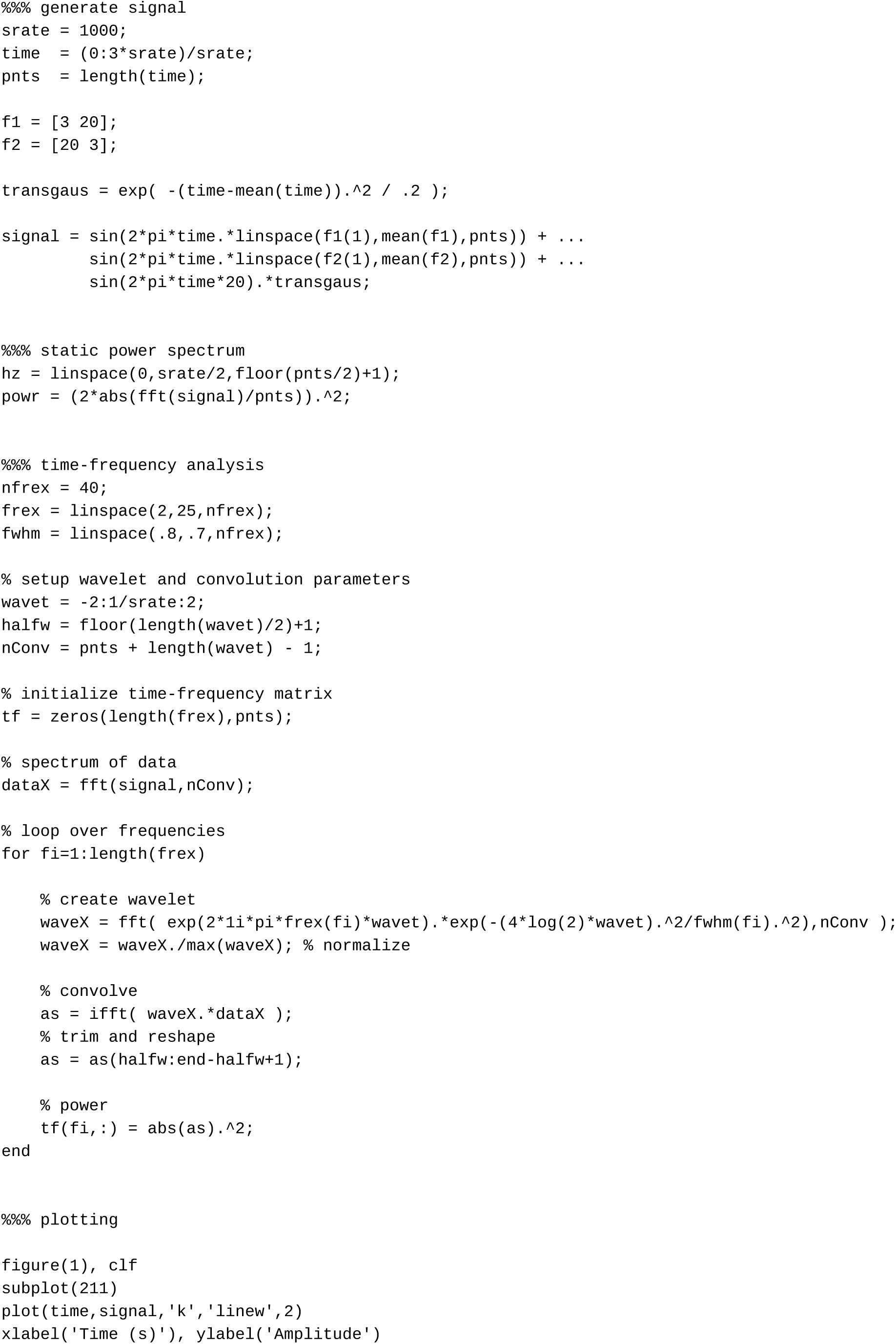
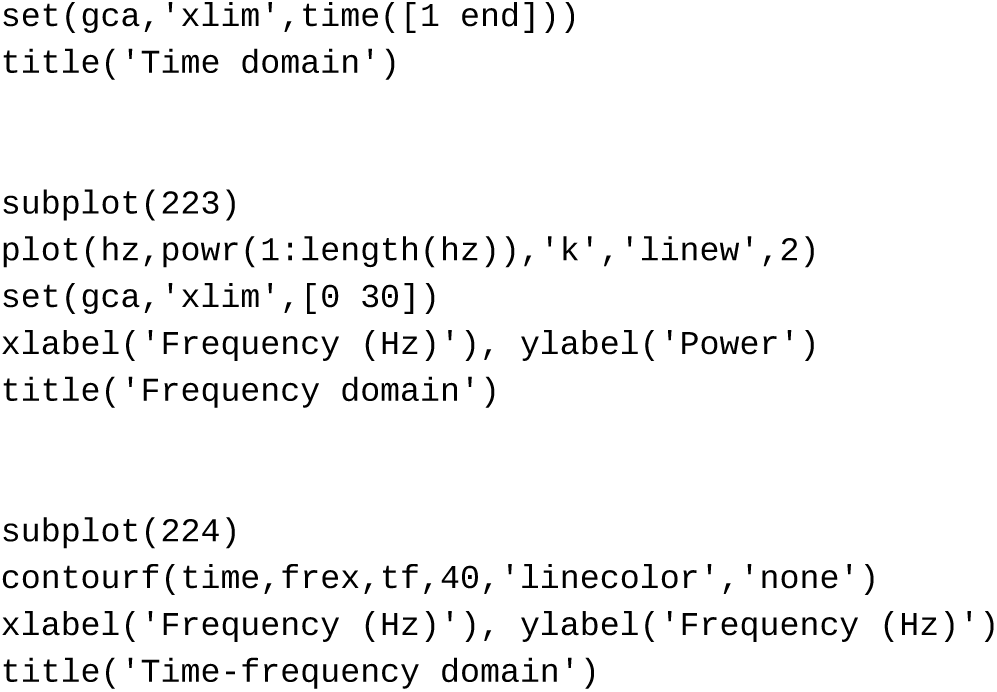

**MATLAB code to reproduce figure 2.**
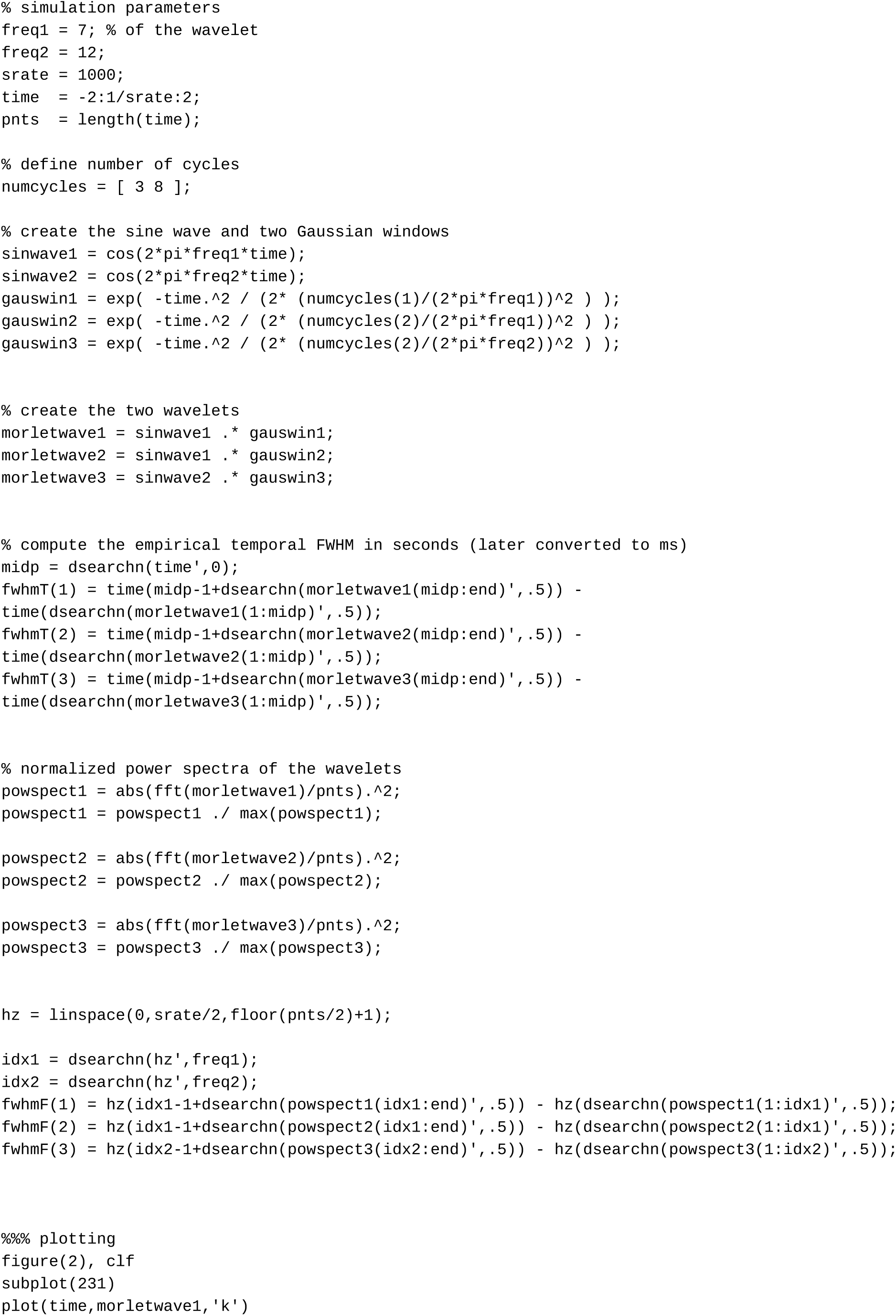
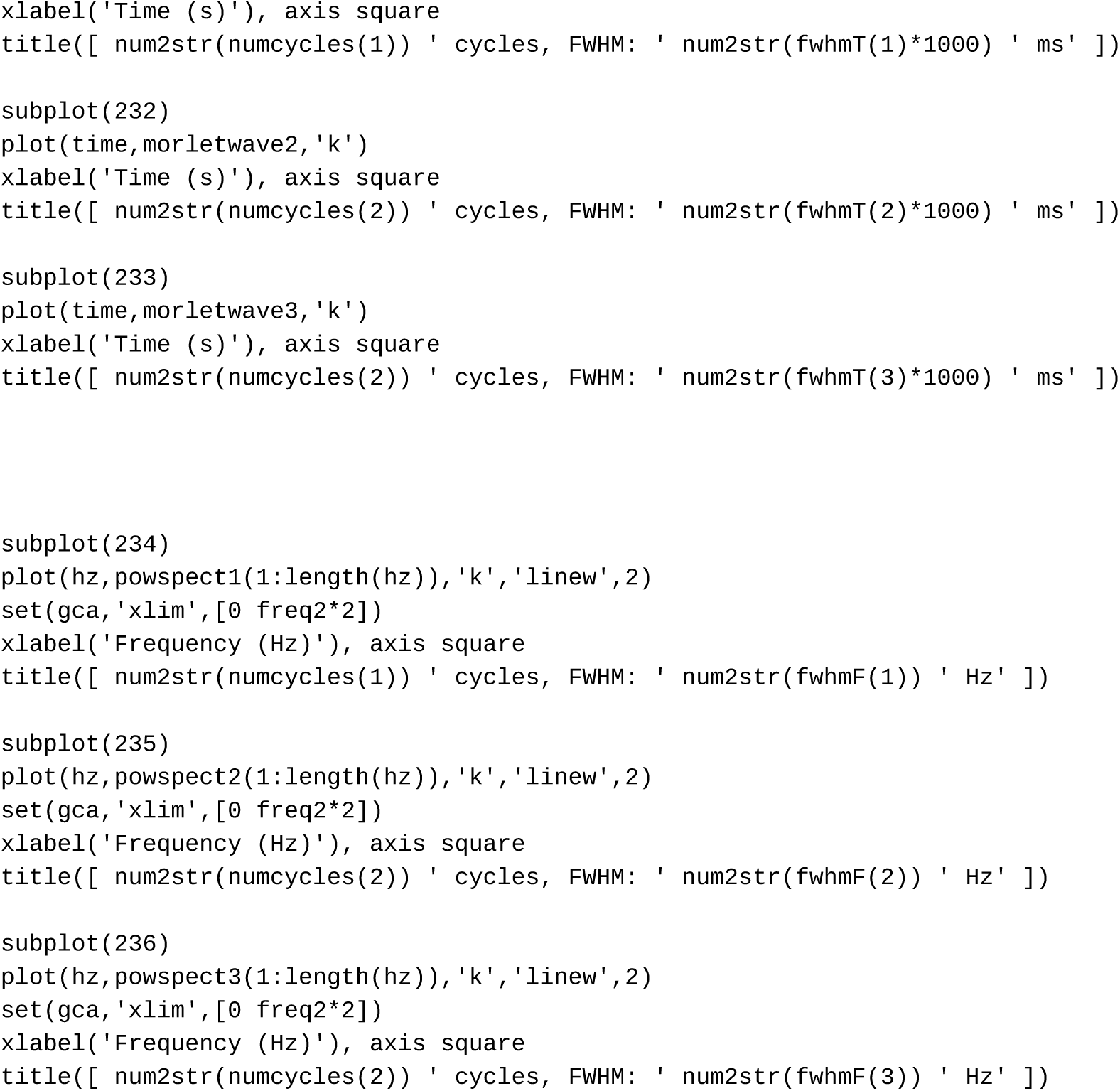

**MATLAB code to reproduce figure 3.**
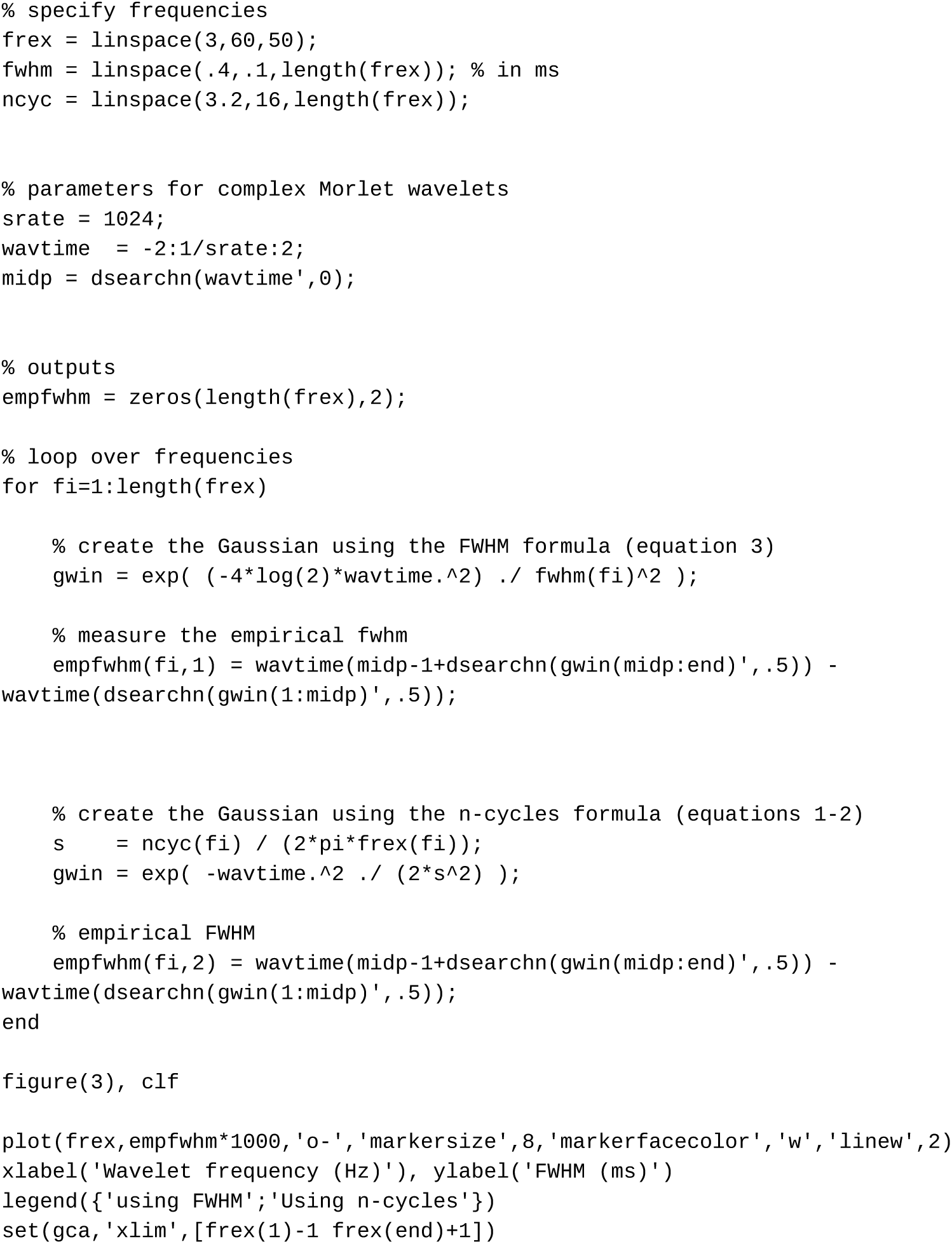

**MATLAB code to reproduce figure 4.**
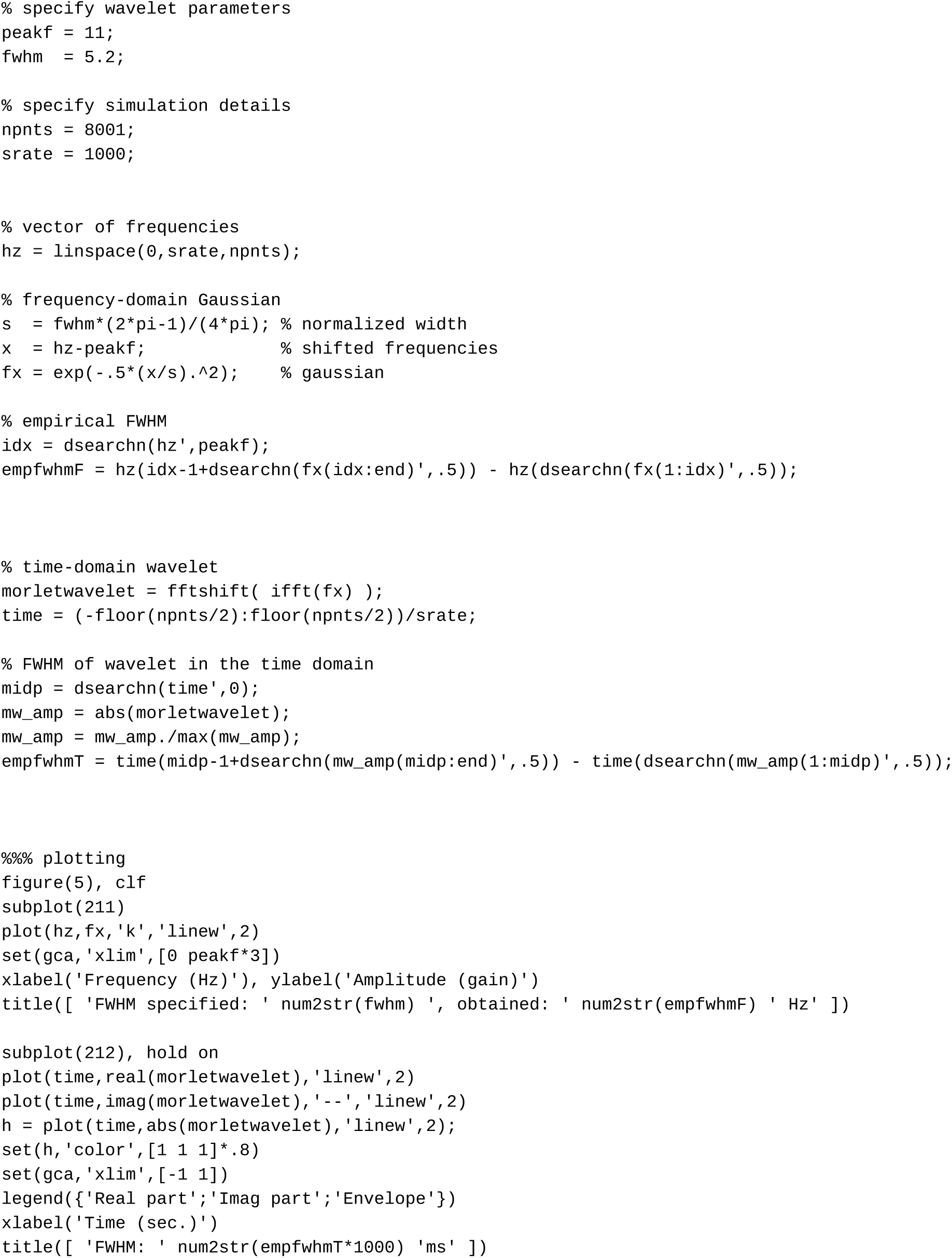

## References

Bruns, Andreas. 2004. “Fourier-, Hilbert- and Wavelet-Based Signal Analysis: Are They Really Different Approaches?” Journal of Neuroscience Methods 137 (2): 321–32.

Cohen, M. X. 2014. Analyzing Neural Time Series Data: Theory and Practice. MIT Press.

Cole, Scott R., and Bradley Voytek. 2017. “Brain Oscillations and the Importance of Waveform Shape.” Trends in Cognitive Sciences 21 (2): 137–49.

Jones, Stephanie R. 2016. “When Brain Rhythms Aren’t ‘rhythmic’: Implication for Their Mechanisms and Meaning.” Current Opinion in Neurobiology 40: 72–80.

